# Colistin heteroresistance in *Enterobacter cloacae* is mediated by PmrAB-independent 4-amino-4-deoxy-L-arabinose addition to lipid A

**DOI:** 10.1101/516872

**Authors:** Katie N. Kang, Dustin R. Klein, Misha I. Kazi, François Guérin, Vincent Cattoir, Jennifer S. Brodbelt, Joseph M. Boll

## Abstract

The *Enterobacter cloacae* complex (ECC) consists of closely-related, but genetically distinct bacteria commonly associated with the human microbiota. ECC have been increasingly isolated from healthcare-associated infections, demonstrating that these Enterobacteriaceae are emerging nosocomial pathogens. ECC strains can rapidly acquire multidrug resistance to conventional antibiotics. Cationic antimicrobial peptides (CAMPs) have served as therapeutic alternatives because they target the highly conserved lipid A component of the Gram-negative outer membrane to lyse the bacterial cell. Many Gram-negative Enterobacteriaceae fortify their outer membrane with cationic amine-containing moieties to protect from CAMP-inflicted lysis. The PmrAB two-component system (TCS) transcriptionally activates 4-amino-4-deoxy-L-arabinose (L-Ara4N) biosynthesis to result in amine moiety addition to lipid A in many Enterobacteriaceae such as *E. coli* and *Salmonella*. In contrast, PmrAB in *E. cloacae* is dispensable for CAMP resistance. Instead, fitness against CAMPs presents as heteroresistance, or a subpopulation of cells that exhibit clinically significant increases in resistance levels compared to the majority population. We demonstrate that *E. cloacae* lipid A is modified with L-Ara4N to induce CAMP heteroresistance and that the regulatory mechanism is independent of the PmrAB_Ecl_ TCS. We show that the response regulator, PhoP_Ecl_, directly binds to the *arnB_Ecl_* promoter to induce expression of L-Ara4N biosynthesis and PmrAB-independent addition to the lipid A disaccharolipid. Therefore, we have identified a mechanism of ECC colistin heteroresistance that directly involves the PhoPQ system.

**Importance:** Members of the *Enterobacter cloacae* complex (ECC) are Gram-negative nosocomial pathogens that have emerged within healthcare facilities around the world. ECC infections are associated with immunocompromised patients and infections are often life threatening. The cationic antimicrobial peptide, colistin (polymyxin E), is a last-line treatment option to combat Gram-negative multidrug resistant infections. However, many ECC intrinsically encode a colistin heteroresistance mechanism. Our analysis to characterize colistin heteroresistance in *E. cloacae* revealed that 4-amino-4-deoxy-L-arabinose is conjugated to the lipid A disaccharolipid to protect from colistin-mediated lysis. Additionally, this mechanism is directly regulated by the PhoPQ_Ecl_ two-component system. Elucidation of outer membrane antimicrobial resistance modifications and their regulatory pathways in *E. cloacae* isolates will advance our understanding of CAMP heteroresistance.

## Introduction

Gram-negative bacteria assemble a highly conserved outer membrane (OM) barrier, which restricts diffusion of toxins such as antibiotics into the cell. Glycerophospholipids comprise the periplasmic monolayer of the asymmetric lipid bilayer, while the surface-exposed monolayer is enriched with lipopolysaccharide (LPS). The LPS glycolipid is organized into three domains; the O-antigen carbohydrate repeat, core oligosaccharide, and the membrane anchor, lipid A (1). The lipid A domain is the bioactive portion of LPS and robustly activates the human Toll-like receptor 4 (TLR-4) and myeloid differentiation factor 2 (MD-2) immune complex to induce an immune response (1–4). Gram-negative pathogens encode highly conserved regulatory mechanisms to promote survival by responding to immune and environmental signals (5–7). Specific signaling pathways regulate lipid A modifications to alter TLR-4/MD-2 recognition and to fortify the OM against immune effectors and antimicrobials, which promotes survival in the host (8).

Lipid A modification enzymes are transcriptionally regulated by two-component systems (TCS) (9, 10). A prototypical TCS consists of an inner membrane sensor histidine kinase (HK) that senses a specific signal and a cognate cytoplasmic response regulator (RR), which alters expression of target genes. Signal recognition induces phosphotransfer and activation of the cognate RR, which typically results in DNA binding to alter gene expression (11). Most HKs encode a phosphatase domain that dephosphorylates the RR when the activating signal is depleted (6, 12). These highly conserved signaling systems enable bacteria to tightly regulate expression of target genes.

The PmrAB and PhoPQ TCSs are well-studied phosphorelay signaling systems that regulate lipid A modifications in response to specific environmental signals (13–15). PmrAB and PhoPQ are highly conserved among pathogenic Enterobacteriaceae, including *Citrobacter*, *Escherichia*, *Klebsiella*, *Salmonella*, and *Shigella* (5). PmrAB responds to high Fe^3+^ concentrations, cationic antimicrobial peptides (CAMPs), and slightly acidic pH to directly activate *eptA* (also known as *pmrC*) and *arn* operon expression (16–18), which encode phosphoethanolamine (pEtN) and 4-amino-4-deoxy-L-arabinose (L-Ara4N) transferases, respectively (19–22). Both enzymes transfer the respective amine-containing chemical moiety onto lipid A at the inner membrane, prior to LPS surface transport (19, 22). Cationic amine addition to the lipid A domain of LPS neutralizes the surface charge to protect the cell from CAMP-mediated lysis (19, 21).

PhoPQ is activated in response to depletion of divalent cations such as Mg^2+^ and Ca^2+^ and the presence of CAMPs (13, 15, 23). PhoPQ phosphotransfer directly activates transcription of genes encoding PagL (only in *Salmonella* (8)) and PagP, which add or remove acyl chains from lipid A, respectively (13, 24–26). While the PmrAB and PhoPQ TCSs each regulate distinct subsets of genes, the independent signaling pathways also converge through crosstalk (27, 28); PmrAB-dependent gene expression is also indirectly regulated by the PhoPQ TCS through the intermediate protein, PmrD. PmrD binds phospho-PmrA to prevent PmrB-mediated dephosphorylation (27, 29–31). Constitutive PmrA-dependent gene expression increases pEtN and L-Ara4N lipid A modifications.

The *Enterobacter cloacae* complex (ECC) is composed of thirteen closely-related Gram-negative bacterial clusters (designated C-I to C-XIII) (32). ECC are typically associated with the host microbiota, but many clusters cause hospital-acquired infections, especially in immunocompromised patients (33). Infections manifest in a wide range of host tissues with symptoms including skin, respiratory tract, urinary tract, wound and blood infections (34). ECC infections have increasingly emerged in nosocomial settings and are problematic because they encode multidrug resistance (MDR) mechanisms, which limits treatment options (33, 35–37). Alternative last-line therapeutics used to treat MDR Gram-negative infections include the CAMP, colistin (polymyxin E), which binds the lipid A portion of LPS to perturb the outer membrane and lyse the bacterial cell. Despite the success of colistin treatment as a last-line therapeutic to combat Gram-negative infections (38, 39), many ECC clusters demonstrated heteroresistance, where a subset of the clonal population is colistin resistant (35, 40–42). We do not fully understand the underlying molecular mechanism(s) that regulate colistin heteroresistance in ECC; further characterization will advance our understanding of antimicrobial resistance and could help improve treatment strategies.

Previous reports showed that colistin heteroresistance naturally occurs within clonal ECC clusters, including *E. cloacae* (cluster XI) (35). Moreover, colistin heteroresistance in *E. cloacae* was induced by innate immune defenses within a murine infection model to lead to treatment failure (40). Transcriptional analysis of susceptible and resistant populations suggested that pEtN and L-Ara4N lipid A modifications contribute to heteroresistance (40) and PhoPQ contributed to regulation (35, 40), as described in other Enterobacteriaceae (5). However, it was not established that the lipid A modifications actually occur, nor has PhoPQ-dependent, PmrAB-independent regulation of colistin heteroresistance been fully described in *E. cloacae* or other ECC isolates.

Herein, we demonstrate that *E. cloacae* colistin heteroresistance is directly regulated by PhoPQ_Ecl_ to induce L-Ara4N modification of lipid A. In contrast, many other Enterobacteriaceae directly regulate L-Ara4N modification of lipid A via the PmrAB TCS. The PhoP_Ecl_ response regulator directly binds to the promoter region of *arnB*_*Ecl*_, which is the first gene of a seven-gene operon (*arnBCADTEF*_*Ecl*_). In contrast, PhoP_Ecl_ does not bind the *arnB* promoter region in *E. coli*. Transcriptomics analysis supports a model of PhoPQ-dependent, PmrAB-independent *arn*_*Ecl*_ regulation. Furthermore, L-Ara4N modification of lipid A increased in response to growth in limiting Mg^2+^, which induced colistin resistance in a PhoPQ_Ecl_-dependent manner.

## Results

### Colistin heteroresistance in *E. cloacae* is regulated by PhoPQ_Ecl_, but not PmrAB_Ecl_

To elucidate the underlying mechanisms that regulate colistin heteroresistance in ECC, we analyzed a collection of *E. cloacae* subsp. *cloacae* strain ATCC 13047 genetic mutants by calculating the colony forming units (CFUs) during exponential growth in the absence and presence of colistin (Fig 1A). While wild type and all mutant *E. cloacae* strains grew in standard growth media, ∆*phoPQ*_*Ecl*_ was not viable when 10 μg/ml of colistin was added to the media. Clinical resistance to colistin is defined as >4 μg/ml (43). The decrease in ∆*phoPQ*_*Ecl*_ cell viability suggested that PhoPQ_Ecl_ signaling is required for colistin heteroresistance. Interestingly, viability was not altered when the ∆*pmrAB*_*Ecl*_ mutant was grown in colistin, which implied that PmrAB_Ecl_ does not regulate colistin heteroresistance. Furthermore, wild type *E. cloacae* grown in colistin demonstrated approximately ten-fold less CFUs at hour two (P value <0.05), suggesting that there was a survival defect in early logarithmic growth phase. However, the fitness defect was no longer significant at hour three and by hour four, CFUs were equivalent to growth without colistin (Fig 1A).

**Figure 1:**
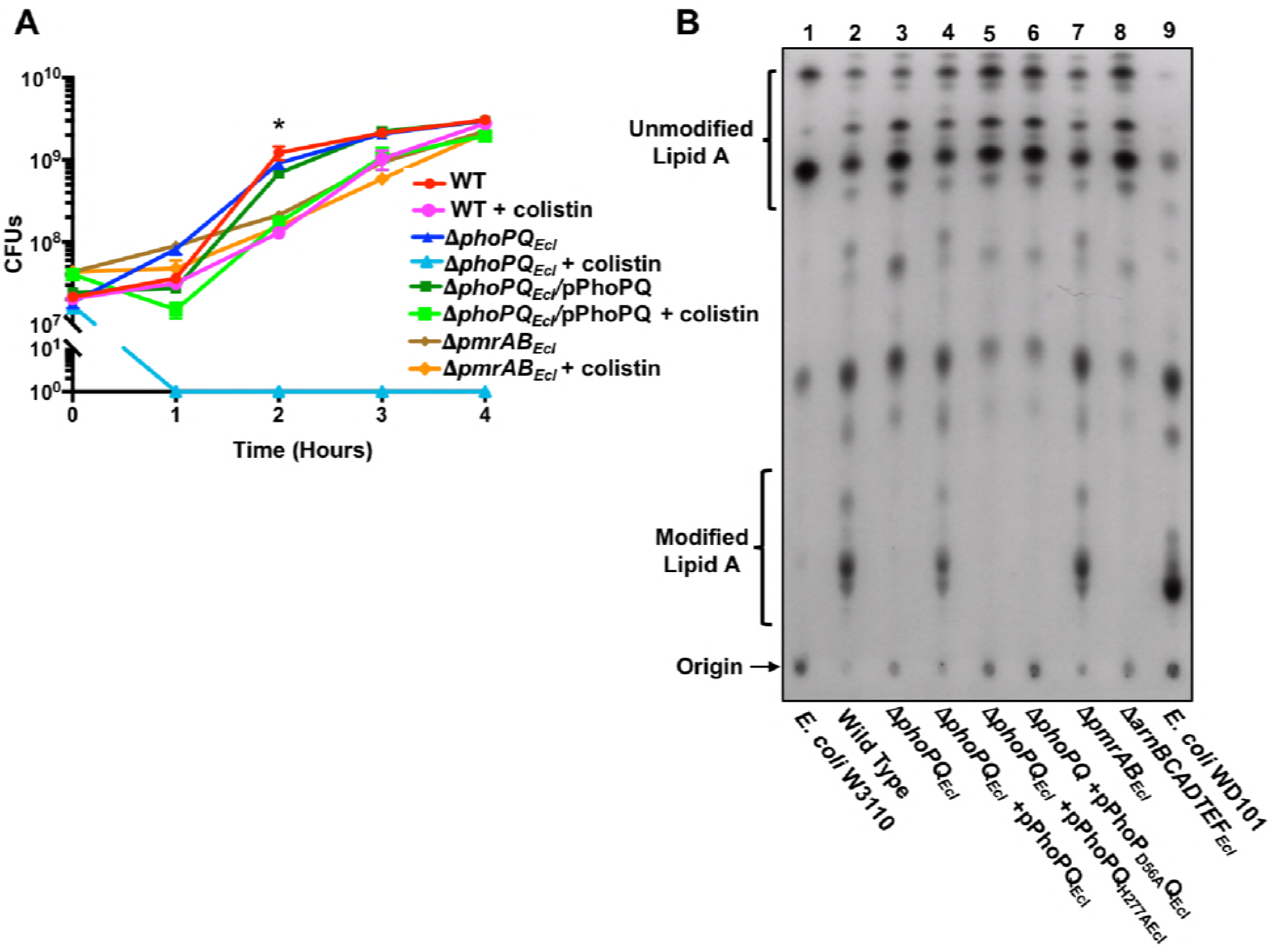
Survival of colistin heteroresistant *E. cloacae* is dependent on PhoPQ, but not PmrAB-regulated lipid A modifications. **(A)** *E. cloacae* logarithmic phase growth over time as measured by colony forming units (CFUs). At two hours, the growth rate between wild type grown in LB was significantly (*) different from cells grown in LB + colistin (P value <0.05). **(B)** ^32^P-radiolabeled lipid A was isolated from wild type and mutant *E. cloacae* strains and separated based on hydrophobicity using thin layer chromatography. Lipid A species are labeled as unmodified or modified as determined by *E. coli* W3110 (lane 1) and WD101 (lane 9) lipid A, respectively.

Due to reports of colistin heteroresistance in *E. cloacae* and other ECC strains (35, 40), we subjected wild type, ∆*phoPQ*_*Ecl*_, ∆*phoPQ*_*Ecl*_/pPhoPQ_Ecl_, and ∆p*mrAB*_*Ecl*_ *E. cloacae* to colistin E-test strip analysis, which provides a convenient method to observe heteroresistance (Fig S1). Squatter colonies within the zone of inhibition indicated colistin heteroresistance in wild type, ∆*phoPQ*_*Ecl*_/pPhoPQ_Ecl_, and ∆p*mrAB*_*Ecl*_ strains, but not ∆*phoPQ*_*Ecl*_. We confirmed colistin heteroresistance using population analysis profiling (PAP) (Table 1). Minimal inhibitory concentration (MIC) values were calculated using the broth microdilution (BMD) method (Table 1). Wild type, ∆*phoPQ*_*Ecl*_/pPhoPQ_Ecl_, and ∆p*mrAB*_*Ecl*_ *E. cloacae* all demonstrated MICs >256 μg/ml, while the ∆*phoPQ*_*Ecl*_, ∆*phoPQ*_*Ecl*_/pPhoPQ_H277A_, ∆*phoPQ*_*Ecl*_/pPhoP_D56A_Q and ∆*arn*_*Ecl*_ (*arnBCADTEF*_*Ecl*_) MIC was 0.5 μg/ml. Together, these studies confirm that PhoPQ_Ecl_ signal transduction and the *arn*_*Ecl*_ biosynthetic operon (L-Ara4N) are required for colistin heteroresistance in *E. cloacae*.

**Table 1.**
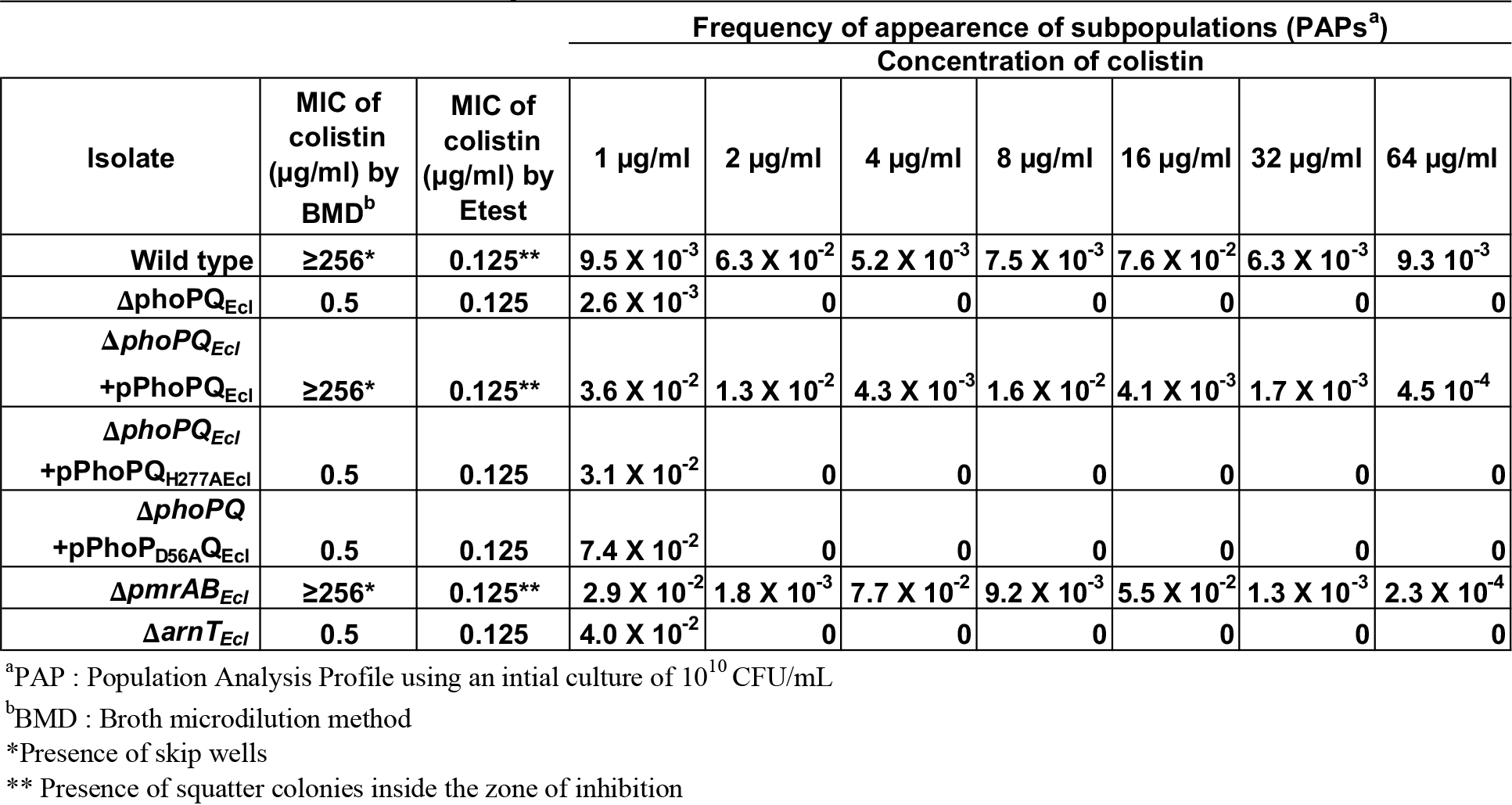
MICs of colistin and PAP analysis for each *E. cloacae* mutant

Since lipid A modifications induce colistin resistance in pathogenic Enterobacteriaceae (5), we analyzed wild type and mutant *E. cloacae* lipid A for modifications. ^32^P-radiolabelled lipid A was isolated and chromatographically separated based on hydrophobicity. As controls, we also analyzed lipid A from *E. coli* strain W3110 (Fig 1B, lane 1), which does not significantly modify its lipid A, and strain WD101(Fig 1B, lane 9), which constitutively expresses *pmrA* to produce modified lipid A (19). Thin layer chromatography (TLC) analysis indicated that wild type *E. cloacae* produced a mixture of lipid A consistent with modified and unmodified species (Fig 1B, lane 2). ∆*phoPQ*_*Ecl*_ and the ∆*arn*_*Ecl*_ strains did not produce modified lipid A (Fig 1B, lanes 3 and 8). PhoPQ complementation fully restored production of modified lipid A in the *phoPQ* mutant (Fig 1B, lane 4). Furthermore, site-directed mutagenesis to substitute H277 in PhoQ_Ecl_ or D57 in PhoP_Ecl_ with alanine limited lipid A assembly to only unmodified species (Fig 1B lanes 5 and 6). These results confirm that PhoPQ_Ecl_ phosphotransfer and L-Ara4N biosynthesis are essential for lipid A modification in *E. cloacae*.

Indirect PhoPQ regulation of lipid A modifications have been described and are conserved among Enterobacteriaceae (5). PhoPQ directly activates PmrD expression, which binds PmrA to induce PmrAB-dependent pEtN and L-Ara4N lipid A modifications (27). In contrast, *E. cloacae* does not encode a PmrD homolog (44). Furthermore, the *pmrAB*_*Ecl*_ mutant assembled a modified lipid A, similar to wild type (Fig 1B, lane 7), and exhibited colistin heteroresistance (Fig 1A, Table 1, Fig S1A), suggesting that PmrAB_Ecl_ does not regulate colistin heteroresistance in *E. cloacae.* Our data contrasts with previous reports describing lipid A modification regulatory pathways in other Enterobacteriaceae, which cannot modify lipid A with L-Ara4N or pEtN when PmrAB signaling is disrupted (19–22, 27).

The lipid A anchor of LPS is a pathogen associated molecular pattern (PAMP) that is bound with high affinity by the mammalian host TLR-4/MD-2 complex (45), which activates a proinflammatory response to clear the bacterial infection (46). Structural alterations to lipid A can dramatically alter TLR-4/MD-2-dependent host immune activation (2) and a previous report nicely demonstrated that *E. cloacae* colistin heteroresistance was induced by innate immune effectors (40). Therefore, we examined if *E. cloacae* containing modified or unmodified lipid A would differentially activate TLR-4/MD-2 in a human embryonic kidney reporter cell line (HEK-blue) (2). Wild type and *phoPQ*_*Ecl*_ mutant strains stimulated TLR-4/MD-2-dependent activation equally (Fig S1B), suggesting that lipid A modifications do not significantly alter host immune recognition. Reporter activation by *E. cloacae* lipid A was attenuated compared to *E. coli* lipid A at higher cell densities, suggesting differential recognition by the human TLR-4/MD-2 complex. The Gram-positive *Staphylococcus aureus* did not produce lipid A and did not stimulate the TLR-4/MD-2 complex (Fig S1B). Thus, while PhoPQ_Ecl_-dependent lipid A modifications contribute to CAMP resistance in *E. cloacae*, they do not significantly affect innate immune recognition and reactivity.

### Determination of *E. cloacae* lipid A modifications

In order to define outer membrane modifications in *E. cloacae*, we isolated lipid A from wild type and ∆*phoPQ*_*Ecl*_. Purified lipid A was analyzed by direct infusion nanoESI. The MS1 spectra with a range of *m/z* 750-2000 are shown in Figure S2. The expanded MS1 spectrum (*m/z* 850-1200) of lipid A isolated from wild type *E. cloacae* demonstrated three distinct modifications: (i) addition of either one or two L-Ara4N moieties (red), (ii) palmitate (C_16:0_) addition (green), and (iii) hydroxylation (Fig 2A) The MS1 spectrum of lipid A isolated from ∆*phoPQ*_*Ecl*_ did not produce L-Ara4N modified lipid A (Fig 2B). Hydroxyl addition was not labeled for simplicity, but correlates with a *m/z* shift of 8 of the doubly-charged molecular ions. Higher-energy collisional dissociation (HCD) and ultraviolet photodissociation (UVPD) MS/MS spectra were obtained for the ions of *m/z* 1042.68 and 1161.79 from wild type and the ions of *m/z* 911.62 and 1030.73 from ∆*phoPQ E. cloacae* (Fig S3, S4, S5 and S6). Analysis of the MS/MS spectra from wild type (*m/z* 1042.68) indicated PhoPQ_Ecl_-dependent addition of L-Ara4N at both the 1- and 4’-phosphates (Fig S3). The MS/MS spectra for the ion of *m/z* 1161.79 (wild type *E. cloacae*) showed addition of L-Ara4N at both the 1- and 4’-phosphates and palmitate addition to the *R*-2-hydroxymyristate (Fig S4). Analysis of lipid A from the *phoPQ*_*Ecl*_ mutant (*m/z* 911.62) completely lacked L-Ara4N modified lipid A (Fig S5) and analysis of the *m/z* 1030.73 ion from the *phoPQ*_*Ecl*_ mutant demonstrated that palmitate addition at the *R*-2-hydroxymyristate position of lipid A occurred independent of PhoPQ_Ecl_ (Fig S6).

Based on transcriptomics studies, a previous report suggested that *E. cloacae* adds pEtN and L-Ara4N to lipid A to develop colistin heteroresistance (40). However, our genetic and high resolution mass spectrometry analysis demonstrate that only L-Ara4N modifies the 1- and 4’-phosphates of lipid A in a PhoPQ_Ecl_-dependent manner (Fig 2A and B) and this amine-containing modification correlates with colistin heteroresistance (Fig 1A and Table 1).

**Figure 2:**
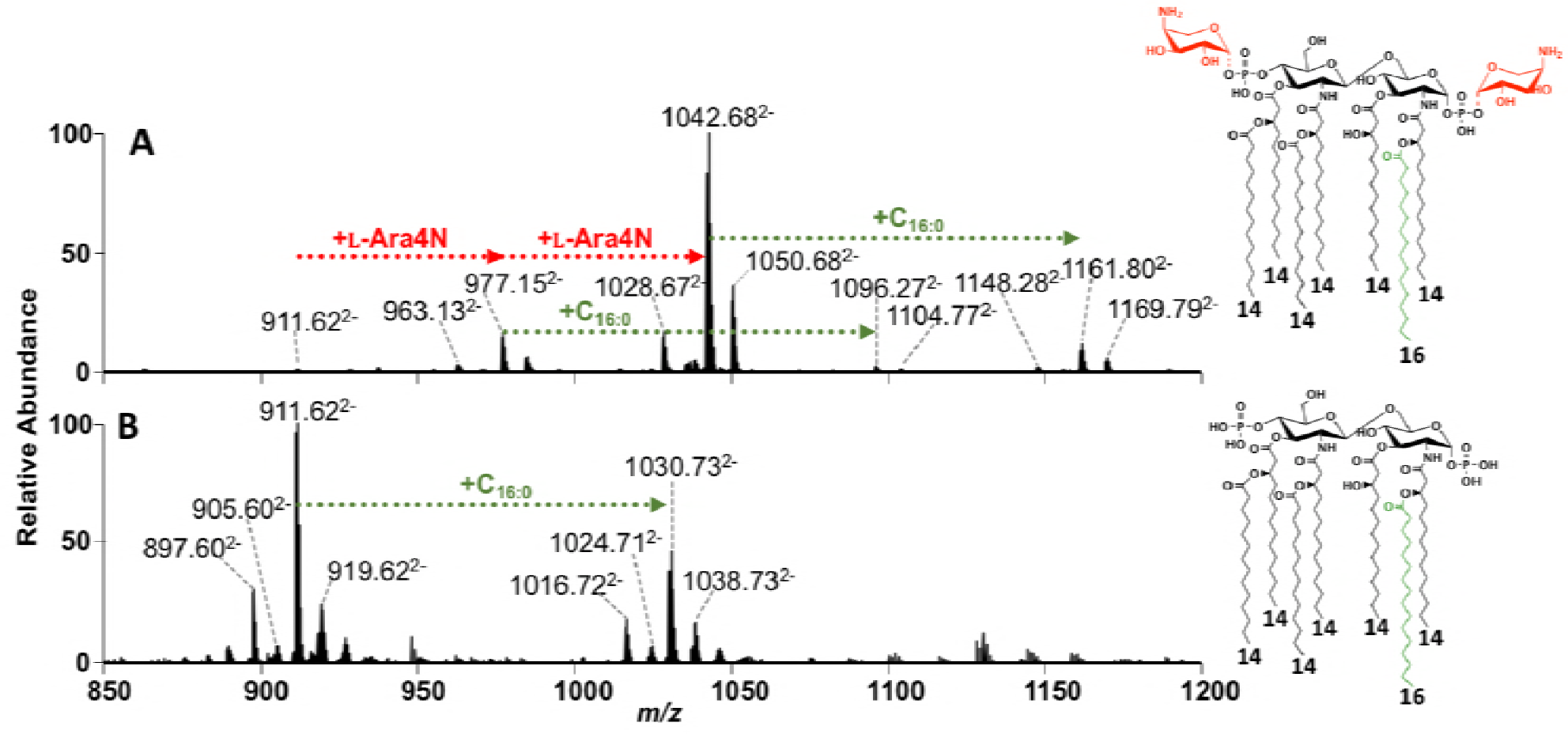
**Expanded MS1 spectra of (A)** Wild type *E. cloacae* lipid A grown with colistin and **(B) Δ***phoPQ*_*Ecl*_ lipid A with the chemical structures associated with the lipid A modifications. The presence of aminoarabinose groups are denoted by L-Ara4N (red), while addition of palmitoyl groups are denoted by +C_16:0_ (green). Hydroxylation is not illustrated, but is indicated by an *m/z* shift of 8 relative to doubly-charged lipid A ions in the spectra.

### L-Ara4N lipid A modifications are dependent on PhoPQ_Ecl_, but not PmrAB_Ecl_

To further characterize lipid A modifications in the ∆*pmrAB*_*Ecl*_ mutant, we analyzed purified lipid A using MALDI-TOF mass spectrometry. While wild type *E. cloacae* produced a lipid A mixture, which included L-Ara4N modified lipids (Fig S7A and B), analysis of ∆*phoPQ*_*Ecl*_ and ∆*arn*_*Ecl*_ indicated that L-Ara4N modified lipid A were not present. Expression of PhoPQ_Ecl_ *in trans* restored L-Ara4N modified lipid A in the *phoPQ*_*Ecl*_ mutant. Importantly, ∆*pmrAB*_*Ecl*_ produced the L-Ara4N modification, similar to wild type (Fig S7A). The *m/z* of each prominent peak in our MALDI-MS analysis corresponded with the exact mass of an expected structure with only the L-Ara4N-containing structures demonstrating colistin resistance (Fig S7B). Here, we confirmed that L-Ara4N modification of lipid A in *E. cloacae* is not dependent on PmrAB_Ecl_.

### PhoP_Ecl_ directly binds to the *arnB*_*Ecl*_ promoter

The *arn* operon is composed of seven genes and expression is driven by a promoter upstream of *arnB* (20). This genetic organization is conserved in *E. cloacae* as illustrated in Fig 3A. *phoP* expression is autoregulated in Enterobacteriaceae, where PhoP binds to the PhoP box where it interacts with RNA polymerase to induce transcription (47). The putative PhoP box in the *phoP* promoter region (P_*phoP*_) is conserved in *E. coli, Salmonella*, and *E. cloacae* (Fig 3B). Alignment of the *E. cloacae arnB* promoter region (P_*arnB*_) with *E. coli, Salmonella*, and *E. cloacae* P_*phoP*_ suggested a putative PhoP box region. Importantly, *E. cloacae* P_*arnB*_, which encodes a putative PhoP box, is highly conserved among ECC. However, this feature was not encoded within *E. coli* P_*arnB*_, suggesting that regulatory mechanisms that control promoter activation are different (Fig 3B).

**Figure 3:**
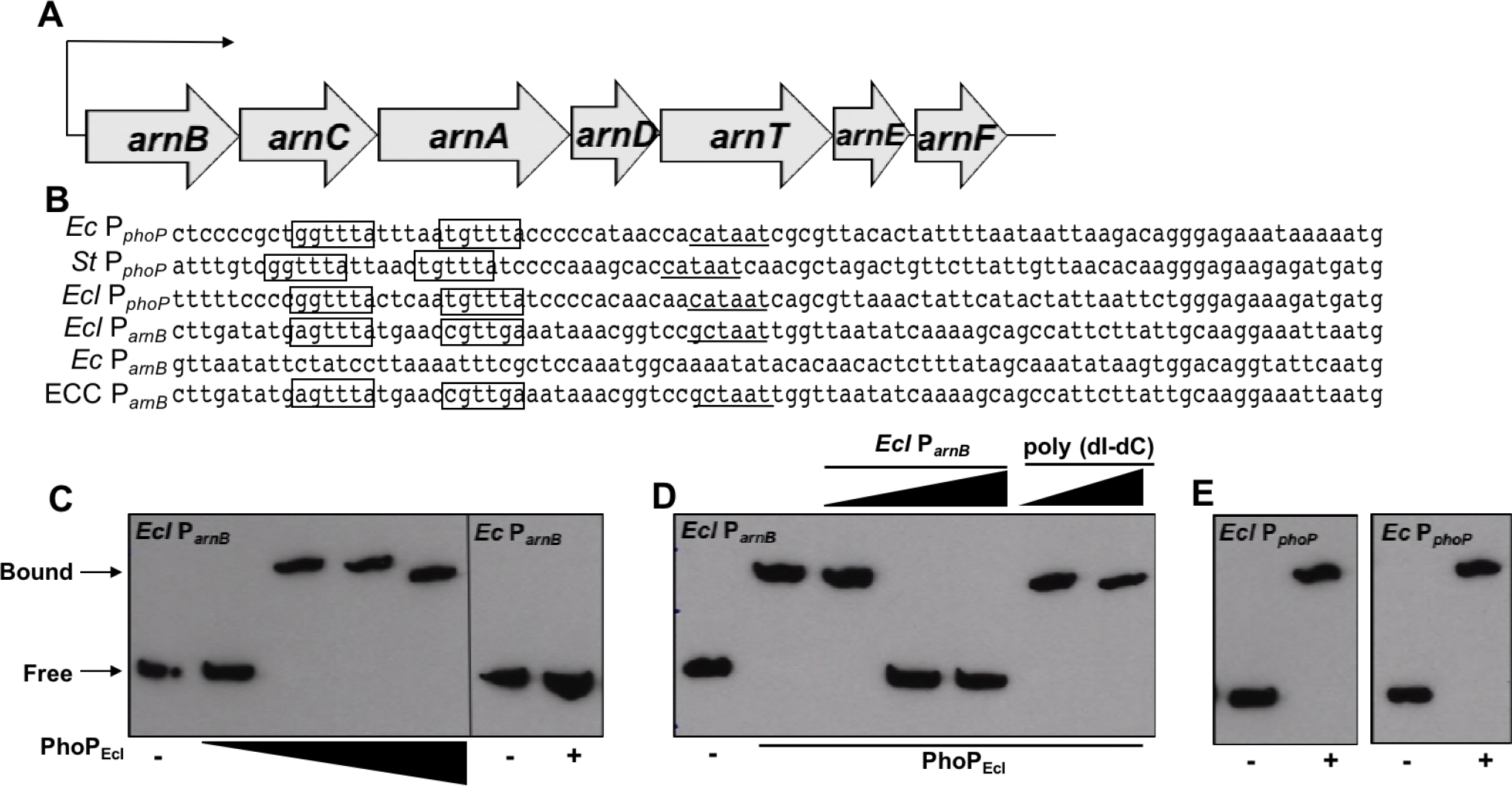
PhoP_Ecl_ binds to the *arnB* promoter of *E. cloacae* (*Ecl*), but not *E. coli* (*Ec*). **(A)** Illustration of the *arn* operon organization. **(B)** Sequence alignment of the *phoP* promoter (P_*phoP*_) region in *Ec*, *Salmonella* (*St*), and *Ecl*, which each contain a PhoP box. The *arnB* promoter (P_*arnB*_) of *Ecl* contains a putative PhoP box binding site that is not present in *Ec.* The putative PhoP boxes have been boxed, while the −10 region is underlined. **(C)** Electrophoretic mobility shift assay (EMSA) of *Ecl* P_*arnB*_ with increasing concentrations of PhoP_Ecl_. PhoP_Ecl_ was used at concentrations of 0, 0.1, 1.0, 5.0 and 10.0 μM. EMSA using *Ec* P_*arnB*_ in the absence or presence of PhoP_*Ecl*_, respectively. **(D)** EMSA competition experiments where increasing concentrations (1:1, 2:1, 5:1) of unlabeled P_*arnB*_ competes with biotin-labeled P_*arnB*_, but nonspecific unlabeled poly (dI-dC) (2:1, 5:1) does not. (E) PhoP_Ecl_ binds to both the *Ecl* and *Ec phoP* promoters.

We performed electrophoretic mobility shifts (EMSAs) using *E. cloacae* P_*arnB*_ to determine if PhoP_Ecl_ directly binds the promoter to activate *arn*_*Ecl*_ transcription. Increasing concentrations of purified PhoP_Ecl_ (Fig S8) induced a shift of the biotinylated *arnB*_*Ecl*_ promoter fragment, which contains the putative PhoP box binding motif (Fig 3C). Importantly, PhoP_Ecl_ does not bind to *E. coli* P_*arnB*_, which does not encode the PhoP box motif (Fig 3C). Furthermore, the PhoP_Ecl_-*arnB*_*Ecl*_ promoter interaction was abrogated when unlabeled *E. cloacae* P_*arnB*_ was added in increasing ratios, as a competitive inhibitor. We also show that the interaction is specific because addition of noncompetitive DNA (poly dI-dC) did not reduce the PhoP_Ecl_ and *E. cloacae* P_*arnB*_ interaction (Fig 3D). Lastly, PhoP_Ecl_ bound *E. cloacae and E. coli* P_*phoP*_, which both encode the nucleotide sequence specific to the PhoP box (Fig 3E). Together, these findings suggest that *E. cloacae* encodes a mechanism that enables L-Ara4N biosynthesis to respond directly to PhoPQ_Ecl_.

### RNA-sequencing analysis of the *phoPQ*_*Ecl*_ and *pmrAB*_*Ecl*_ mutants

To better understand the PhoP_Ecl_ and PmrAB_Ecl_ regulatory products, we isolated and sequenced total RNA from wild type and mutant *E. cloacae* strains. A heat map illustrates the fold change of *arn*_*Ecl*_, *phoPQ*_*Ecl*_, and *pmrAB*_*Ecl*_ gene expression in the TCS mutants relative to wild type (Fig 4). Expression of the *arn*_*Ecl*_ genes were significantly down regulated in ∆*phoPQ*_*Ecl*_ compared to wild type, suggesting that activation of the pathway is dependent on PhoPQ_Ecl_. In contrast, *arn*_*Ecl*_ gene expression was not significantly altered in the ∆*pmrAB*_*Ecl*_ mutant relative to wild type.

**Figure 4:**
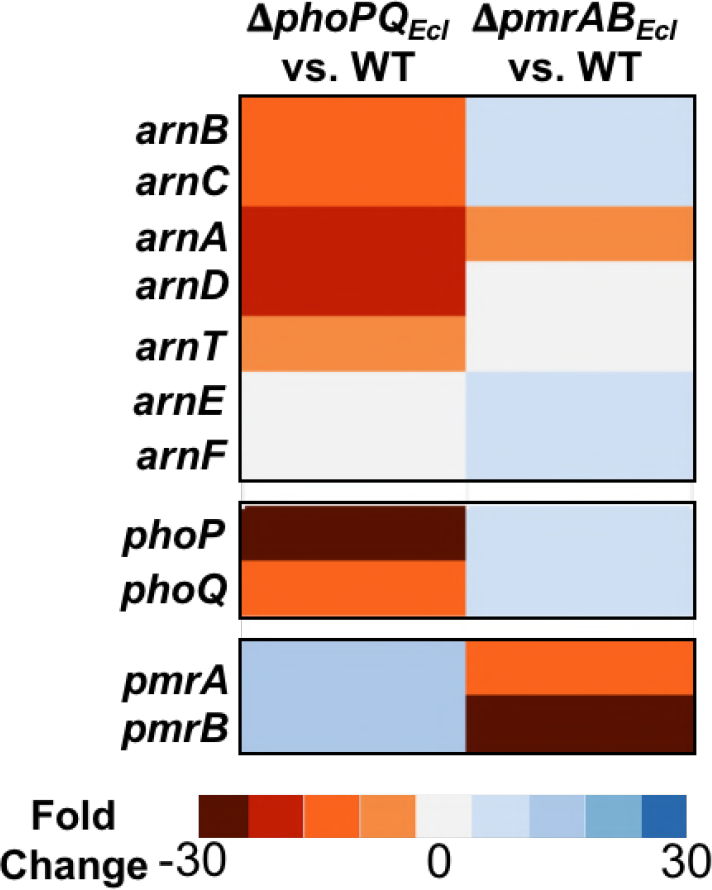
RNA-sequencing analysis of *E. cloacae genes*. Heat map illustrating the altered expression of select operons in **Δ***phoPQ*_*Ecl*_ and **Δ***pmrAB*_*Ecl*_ mutants. Expression is shown as a ratio of mutant to wild type expression (P <0.05).

### Colistin resistance increases within the *E. cloacae* population in response to limiting Mg^2+^

Together the transcriptomic analysis (Fig 4) and colistin heteroresistance in wild type *E. cloacae* (Fig 1A and Table 1) suggest constitutive addition of L-Ara4N to lipid A in a subset of the population under standard growth conditions. In *E. coli* and *Salmonella*, PhoPQ is activated by various signals, including low Mg^2+^ and CAMPs (13, 23, 25). PhoP activates PmrAB, which stimulates pEtN and L-Ara4N addition to lipid A (27). Here we analyzed if PhoPQ_Ecl_ responds to similar physiological cues to induce colistin resistance in *E. cloacae*. Wild type and mutant *E. cloacae* were grown in N minimal medium with high (10 mM) or low (10 μM) Mg^2+^ levels. All cultures were exposed to colistin at mid-logarithmic growth. Wild type and complemented *phoPQ*_*Ecl*_ mutant strains grown in high Mg^2+^ demonstrated some susceptibility to 5 and 10 μg/ml of colistin (Fig 5A, High Mg^2+^), suggesting colistin-susceptible and-resistant populations were present. When grown under limiting Mg^2+^ conditions, *E. cloacae* cells were resistant (Fig 5A, Low Mg^2+^). The *phoPQ*_*Ecl*_ mutant demonstrated a fitness defect in either Mg^2+^ concentration when exposed to colistin (Fig 5A). These data suggest that PhoPQ_Ecl_ induces colistin resistance in response to limiting Mg^2+^ growth conditions.

**Figure 5:**
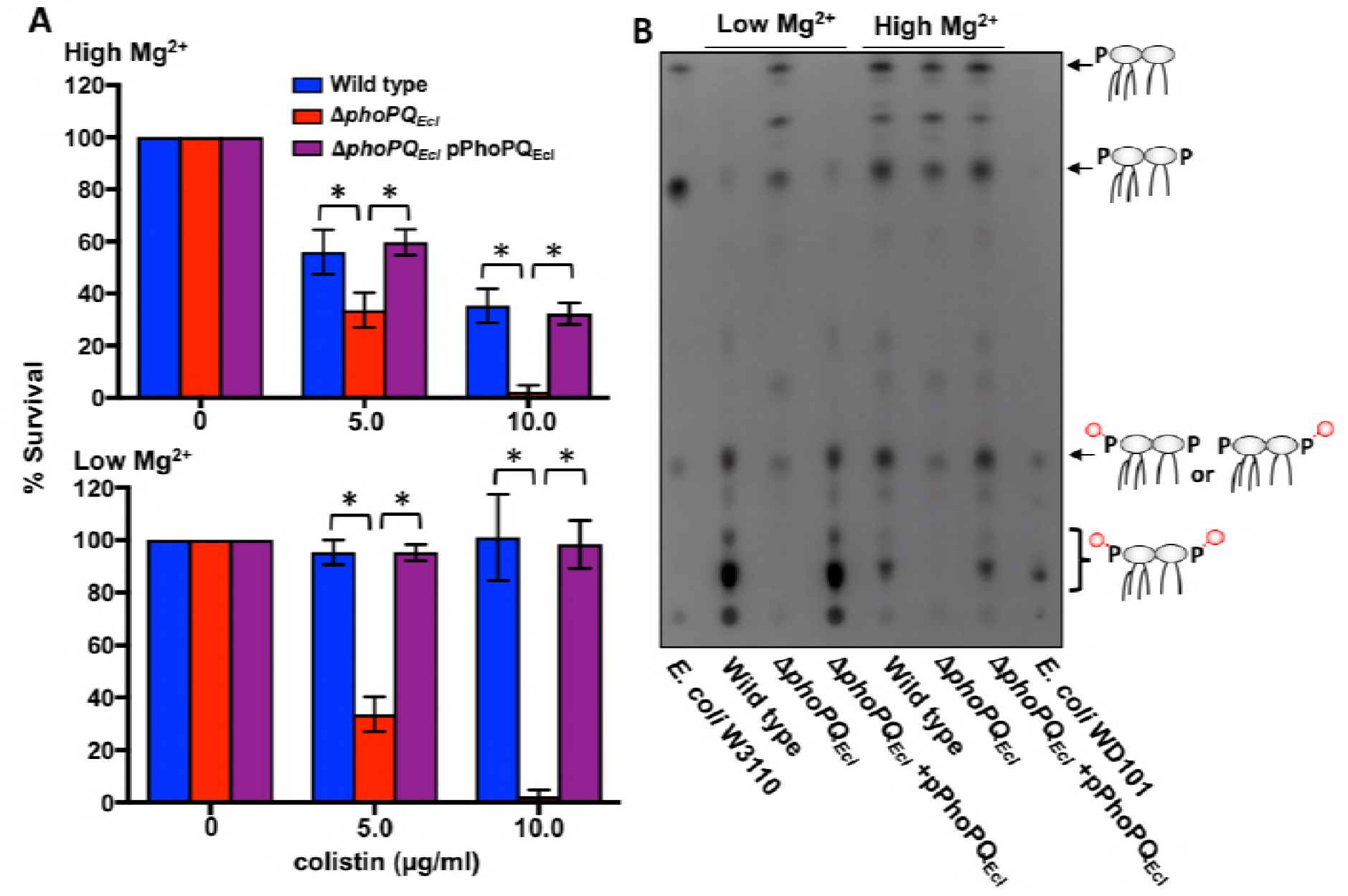
PhoPQ_Ecl_-dependent activation of l-Ara4N addition induces colistin resistance in low Mg^2+^. **(A)** Wild type and mutant *E. cloacae* strains were grown in N minimal medium with low (10 μM, top) or high (10mM, bottom) Mg^2+^. Strains were challenged with 0, 5, or 10 μg/ml of colistin for 1 h and plated for survival. Two biological replicates were each analyzed in triplicate with data from one representative set reported. *P* value <0.05. **(B)** ^32^P-radiolabeled lipid A was isolated from wild type and mutant *E. cloacae* strains and separated based on hydrophobicity using thin layer chromatography. The associated lipid A structures are illustrated with red circles indicating L-Ara4N Addition. Lipid A species were labeled as determined by *E. coli* W3110 (unmodified) and WD101 (modified) lipid A.

### PhoPQ_Ecl_ responds to limiting Mg^2+^ conditions by inducing L-Ara4N lipid A modification

To determine if increased colistin resistance was dependent on L-Ara4N modification of lipid A, we isolated lipid A after growth in either low or high Mg^2+^. TLC analysis demonstrated that wild type and the complemented *phoPQ*_*Ecl*_ mutant primarily produced L-Ara4N-modified lipid A when Mg^2+^ concentrations were limiting (Fig 5B, Low Mg^2+^). In contrast, the same strains grown in excess Mg^2+^, produced a mixture of modified and unmodified lipid A (Fig 5B, High Mg^2+^). Interestingly, growth in excess Mg^2+^ does not completely shut-off production of PhoPQ_Ecl_-dependent lipid A modification in *E. cloacae*, as was previously shown in *E. coli* (27). We did observe a decrease in the relative amount of the di-L-Ara4N form while increasing the ratio of unmodified lipid A species (Fig 5B). Together, these studies suggest that a subset of the clonal *E. cloacae* population activates PhoPQ_Ecl_-dependent L-Ara4N of lipid A under standard growth conditions. However, depletion of Mg^2+^ induces PhoPQ_Ecl_ signaling to enhance L-Ara4N modification (Fig 5B) and colistin resistance (Fig 5A) throughout the population.

## Discussion

*E. cloacae* and other ECC members encode PmrAB_Ecl_ and PhoPQ_Ecl_ homologs, which we hypothesized would function in a signaling pathway to regulate L-Ara4N and pEtN modification of lipid A based on previous transcriptomics analysis of resistant and susceptible populations (40) and because these lipid A modifications are highly conserved among Enterobacteriaceae (5). However, our genetic and high-resolution mass spectrometry analysis of *E. cloacae* lipid A determined that colistin heteroresistance *in E. cloacae* was mediated by PhoPQ_Ecl_-dependent L-Ara4N lipid A modification. Therefore, we have identified a mechanism of ECC colistin heteroresistance that involves the PhoPQ system.

*E. cloacae* and other ECC members do not encode a PmrD homolog, which couples PhoPQ signal transduction to regulation of PmrA-dependent genes in many Enterobacteriaceae (5). Moreover, PmrA_Ecl_ shares only 52% identity with *E. coli* PmrA and PmrB_Ecl_ shares only 57% identity with *E. coli* PmrB, suggesting the L-Ara4N lipid A modification pathway in *E. cloacae* diverged from other Enterobacteriaceae. We confirmed direct binding of PhoP_Ecl_ to the *arnB*_*Ecl*_ promoter, which supports a model where L-Ara4N addition to lipid A and colistin heteroresistance in *E. cloacae* is dependent on PhoPQ_Ecl_, but not PmrAB_Ecl_.

Research from other groups has outlined a complex regulatory network in *E. coli* and *Salmonella* that tightly regulates lipid A L-Ara4N and EptA modifications (19–22, 27). We hypothesize that uncoupling PmrAB_Ecl_ signal transduction from L-Ara4N modification bypasses a key regulatory checkpoint, which likely promotes constitutive *arn*_*Ecl*_ transcription and L-Ara4N modification of lipid A in a subset of the clonal *E. cloacae* population. Since selection has driven *E. cloacae* and other ECC to maintain an altered lipid A modification signaling network, we predict that it is advantageous to maintain a CAMP resistant subpopulation in some environments. Presumably, the alternative regulatory mechanism promotes bacterial fitness in environments specific to its commensal and pathogenic niches.

While the dynamics of the regulatory cascades that control lipid A modification in Gram-negative bacteria have been extensively characterized in *E. coli* and *Salmonella* and generalized across Enterobacteriaceae, mechanisms that regulate lipid A modifications in *E. cloacae* highlight variations that promote clinically important resistance levels. Furthermore, colistin heteroresistance has also been associated with *Klebsiella pneumoniae* (48), another Enterobacteriaceae family member, which highlights the importance of studying antimicrobial resistance mechanisms at the species and strain level.

## Materials and Methods

### Bacterial Strains and Growth

*E. cloacae* subsp. *cloacae* ATCC 13047 and ECC strains were initially grown from freezer stocks on Luria-Bertani (LB) agar. Isolated colonies were used to inoculate LB broth or N minimal medium (0.1M Bis-Tris, pH 7.5 or 5.8, 5 mM KCl, 7.5 mM (NH_4_)_2_SO_4_, 0.5 M K_2_SO_4_, 1 mM KH_2_PO_4_, 0.10% casamino acids 0.2% glucose, 0.0002% thiamine, 15 μM FeSO_4_, 10 μM or 10 mM MgSO_4_) at 37° C. Kanamycin was used at 25 μg/ml for selection and colistin was used at 5 μg/ml or 10 μg/ml where indicated.

All strains and plasmids used in this study are listed in Table S1. Briefly, *E. cloacae* subsp. *cloacae* 13047 mutant strains were constructed as previously described using recombineering with the plasmid pKOBEG (49). Linear PCR products were introduced in to the *E. cloacae* ATCC 13047/pKOBEG strain by electroporation and plated on selective media. Selected clones were transformed with pCP20 to cure the antibiotic resistance cassette.

To complement *E. cloacae* mutants, the coding sequence from *phoPQ*_*Ecl*_ was cloned into the SalI and KpnI sites in pMMBKn (3). To generate point mutants in PhoQ_H277A_ and PhoQ_D56A_, site directed mutagenesis was performed using Pfu Turbo using primers that incorporated the associated alanine-encoded nucleotide replacement. All constructs were validated using Sanger sequencing. IPTG inducible constructs were transformed into the *phoPQ* mutant and grown in 2.0 mM IPTG to induce expression.

### Colony Forming Unit Counts

*E. cloacae* subsp. *cloacae* 13047 and mutant strains were initially grown from freezer stocks on Luria-Bertani (LB) agar. Isolated colonies were resuspended and used to inoculate LB broth with 10 μg/ml or without colistin at an OD_600_ = 0.01. Cells were plated at designated time points on LB agar. Plates were grown overnight at 37° C and colony forming units (CFU) were counted and reported.

### Broth Microdilution assays

MICs of colistin were determined in triplicate by the broth microdilution (BMD) method. Briefly strains were inoculated from overnight cultures at an OD_600_ = 0.1. Various concentrations (0 - 256 μg/ml) of colistin were added to each well and cultures were incubated overnight. Growth was indicated by a reading the OD_600_ and the lowest concentration at which growth was inhibited was recorded as the MIC. *E. coli* W3110 and WD101 were used as control strains. In some cases, ‘skip wells’ were observed suggesting a heteroresistance phenomenon and the MIC was determined disregarding the clear wells (35).

### Population Analysis

Population analysis profiling was performed by plating a high inoculum (1 X 10^10^ CFU) onto LB agar containing 1 to 64 μg/ml colistin (in 2-fold increments). Plates were incubated overnight at 37° C and frequency of the subpopulation was determined by dividing by the total number of cells (50).

### Isolation of Lipid A

Isolation of lipid A for TLC analysis involved ^32^P-radiolabeling of whole cells was performed as previously described with slight modifications (51). In brief, 12.5 ml of *E. cloacae* was grown at 37° C to OD_600_ = 1.0. Bacteria were harvested by centrifugation at 10,000 X g for 10 min. Lipid A extraction was carried out by mild-acid hydrolysis as previously described (52).

### Mass Spectrometry

MS1 spectra of lipid A in Figure 4 were collected on a MALDI-TOF/TOF (Axima Performance, Shimadzu) mass spectrometer in the negative mode. All other spectra were collected in the negative mode on a Thermo Scientific Orbitrap Fusion Lumos mass spectrometer (San Jose, CA, USA) modified with a Coherent ExciStar XS ArF excimer laser (Santa Clara, CA), as previously described (53). HCD was performed with the normalized collision energy (NCE) of 25%. UVPD was performed with the laser emitting 193 nm photons at 5 mJ per laser pulse with 5 pulses per scan. The laser pulse repetition rate was 500 Hz. The instrument was operated at 120000 resolving power with a precursor isolation window of 3 *m/z*. All samples were dissolved in 50:50 MeOH:CHCl_3_ and directly infused into the mass spectrometer via a static nano-electrospray ionization source. The presented spectra are an average of 50 scans.

### TLR-4 Signaling Assays

HEK-Blue hTLR4, cell line was maintained according to the manufacturer specifications (Invivogen). Overnight bacterial cultures in stationary phase were serial diluted for assays as previously described (2, 3). At least two biological replicates were each done in triplicate and one representative set was shown.

### Colistin Survival Assays

Polymyxin E survival assay analysis were performed as previously described with slight modifications (27). Wild type and mutant *E. cloacae* strains were grown overnight on LB agar. The following day, media N minimal medium with pH = 7.5, 10 μM MgSO_4_ were inoculated at OD_600_ = 0.1 with bacteria from overnight cultures that were washed with N minimal media without Mg^2+^ or iron. Cultures were grown until OD_600_ = 0.6, when they were split and treated with 0, 5 or 10 μg/ml of colistin (Polymyxin E). Cultures were incubated for 1 h at 37° C and then colony-forming units were plated and calculated. Percent survival was calculated by dividing the number of bacteria after treatment with colistin relative to those incubated in the absence of colistin and then multiplied by 100.

### Protein Purification

To purify the PhoP_Ecl_ protein, the coding sequence was cloned into pT7-7Kn, as previously described (7). Briefly the *phoP*_*Ecl*_ CDS was amplified from *E. cloacae* cDNA with primers that added a C-terminal His_8X_ tag. From an overnight starter culture, 1 Liter of LB broth containing 25 μg/ml of kanamycin was inoculated at 1:50 and grown at 37° C until the OD_600_ = 0.5. IPTG was added to a final concentration of 1mM, and the culture was incubated at 37° C for an additional 4 h. Bacteria were recovered by centrifugation at 10,000 × g for 10 min, and the bacteria were resuspended in lysis buffer. Bacteria were lysed using sonication and the soluble fraction was recovered by centrifugation at 10,000 × g for 30 min. PhoP_Ecl_-His_8X_ was purified on a Ni-nitrilotriacetic acid (NTA) beads according to the manufactures instructions (Qiagen).

### Electrophoretic Mobility Shift Assay

PhoP_Ecl_-His_8X_ proteins were purified as described above. EMSAs were performed based on a modified protocol (6). 250-bp DNA fragments of *phoP*_*Ecl*_ and *arnB*_*Ecl*_ spanning −230 to +20 relative to the translational start site were amplified from *E. cloacae* or *E. coli* cDNA using 5’-biotinylated primers. PhoP_Ecl_-His_8X_ proteins were incubated with biotinylated DNA at 25° C for 20 min. For competition experiments, unlabeled *E. cloacae* P_*arnB*_ and poly (dI-dC) were added at 1:1, 2:1, or 5:1 ratios relative to biotin-labeled P_*arnB*_ DNA, and 0.1 - 10 μM of PhoP_Ecl_-His_8X_ proteins were used. After electrophoresis at 4° C, protein/DNA was transferred onto a positively charged nylon membrane. Blots were blocked in 5% milk in TBS for 20 min and streptavidin conjugated HRP was used at a 1:300 dilution.

### Nucleic Acid Extraction

Total RNA was extracted using the Direct-Zol RNA MiniPrep Kit (Zymo Research) from *E. cloacae* grown to a final OD_600_ = 0.6. Isolated RNA was treated with DNA-free DNA removal kit (Thermo-Fisher Scientific) to eliminate genomic DNA contamination. DNase-depleted RNA was used for qRT-PCR and RNA-seq.

### RNA-sequencing

RNA-sequencing was performed as previously described (54). Briefly, DNA-depleted RNA was processed for Illumina sequencing using the NEB Next Ultra Directional RNA Library Prep kit for Illumina as described by the manufacturer (NEB). Sequencing was performed using Illumina HiSeq. Sequencing data was aligned to the *E. cloacae* subs. *cloacae* ATCC 13047 published genome annotations (44) using CLC genomic workbench software (Qiagen) and RPKM expression values were determined. The weighted proportions fold change of expression values between samples was determined and a Baggerley’s test on proportions was used to generate a false discovery rate corrected P-value. We then used a cut-off of 2-fold weighted proportions absolute change with a false-discovery rate corrected P-value of ≤ 0.05 to identify significantly differentially regulated genes between samples. The sequencing data for the clinical isolates has been deposited in the Nation Center for Biotechnology’s Gene Expression Omnibus (PRJNA461875).

## Acknowledgements

This work was supported by NIH GM103655 (J.S.B), and Welch Foundation F-1155 (J.S.B.). Funding from the UT System for support of the UT System Proteomics Core Facility Network is gratefully acknowledged.

We would like to thank Cara Boutte and Mark Pellegrino for thoughtful review of the manuscript.

